# The control and training of single motor units in isometric tasks are constrained by a common synaptic input signal

**DOI:** 10.1101/2021.08.03.454908

**Authors:** Mario Bräcklein, Jaime Ibáñez, Deren Y Barsakcioglu, Jonathan Eden, Etienne Burdet, Carsten Mehring, Dario Farina

## Abstract

Recent developments in neural interfaces enable the real-time and non-invasive tracking of motor neuron spiking activity. Such novel interfaces provide a promising basis for human motor augmentation by extracting potential high-dimensional control signals directly from the human nervous system. However, it is unclear how flexibly humans can control the activity of individual motor neurones to effectively increase the number of degrees-of-freedom available to coordinate multiple effectors simultaneously. Here, we provided human subjects (N=7) with real-time feedback on the discharge patterns of pairs of motor units (MUs) innervating a single muscle (tibialis anterior) and encouraged them to independently control the MUs by tracking targets in a 2D space. Subjects learned control strategies to achieve the target-tracking task for various combinations of MUs. These strategies rarely corresponded to a volitional control of independent input signals to individual MUs. Conversely, MU activation was consistent with a common input to the MU pair, while individual activation of the MUs in the pair was predominantly achieved by alterations in de-recruitment order that could be explained with history-dependent changes in motor neuron excitability. These results suggest that flexible MU control based on independent synaptic inputs to single MUs is not a simple to learn control strategy.

## 1. Introduction

Decoding single motor unit (MU) spiking activity, i.e. action potentials discharged by motor neurons and their innervated muscle fibres, non-invasively from the surface electromyogram (EMG), represents a viable alternative to invasive brain recordings for neural human-machine interfaces [1]–[3]. One potential application for such non-invasive neural interfaces is to augment the number of degrees-of-freedom a person can control by exploiting the fact that hundreds to thousands of motor neurons innervate muscles [4]. If single MUs could be individually controlled, they could potentially be activated in a multitude of ways, leading to an enormous potential for additional information transfer. New patterns made of independent MU control could then be used to provide the basis for augmented control signals without impeding the original function of the innervated muscles for natural limb control [5], [6]. In fact, the force a muscle exerts reflects the common activity of populations of MUs [7]. In contrast, the independent control of some individual MUs would result in a high-dimensional activation without any major impact on the force, in the same way as the activity in single MUs determined by independent synaptic noise is filtered in force production [8]. Therefore, the possibility of controlling part of the MUs in a muscle independently would indeed provide a separation from the natural control of force, mainly provided by the population behaviour, and a vast augmentation resource from independent control. Nonetheless, while being a very attractive potential mechanism for artificial augmentation, the independent control of MUs would imply an increased computational load by the central nervous system (CNS) without any known *natural* functional benefit for the human motor system. For this reason, most previous observations indicate that, contrary to independent control, single MUs tend to be activated in a very stereotyped way which is determined by the common input received by functional groups of MUs [8]–[10] and by their biophysical properties (i.e., *Henneman’s size principle* stating a dependency between the exerted force at which a MU starts to contribute and the neuron’s size [11], [12]). Since the size of the soma is inversely related to the membrane resistance, smaller motor neurons discharge action potentials earlier and faster than larger neurons for the same net excitatory synaptic input [13], [14]. For this reason, if a pool of MUs receives the same common input, MU recruitment is solely dependent on the MU anatomy and on the intrinsic excitability of the motor neurones. The size principle has been observed in several muscles [15]–[20] and appears to remain robust in various scenarios [21], [22].

Previous works have tried to challenge the perspective of single MUs only being activated in a pre-determined fashion [2], [23]. A recent study even provided evidence that indeed there is, to some degree, a neural substrate that would allow for the selective cortical control of MUs via descending pathways [24]. However, so far, it is unclear whether humans can learn to leverage such a potential neural structure for selectively activating MUs by converting the common neural input received by a MU pool to independent inputs to individual MUs and thus change the original MU recruitment.

This study examined whether humans could control pairs of MUs innervating the same muscle flexibly. Further, it addressed how this potential ability depends on the similarity of the MU pairs in size or, equivalently, in recruitment threshold. For this purpose, we used a neural interface that provided subjects with biofeedback on the activity of individual MUs [25]. Subjects were encouraged to navigate a cursor inside a 2D space into different targets as quickly as possible by selectively recruiting different MUs. This allowed us to assess if subjects were able to leverage potential selective descending pathways that would facilitate independent synaptic input to individual MUs or if, instead, they used control strategies based on a common input to the MU pair. After several days of training, all subjects achieved the target-tracking task. However, the control strategies used that allowed individual MU activation did not leverage on potential selective inputs to single MUs. Instead, subjects strongly favoured control strategies based on a common input signal combined with changes in intrinsic motor neuron excitability due to history-dependent physiological properties of the activated MUs.

## 2. Methods

### 2.1. Subjects

Seven healthy subjects (two females and five males, age: 27.86 ± 4.06 years [μ ± SD]) were recruited for the study of whom three are authors of this article. Four subjects were naïve to the experimental paradigm, while the remaining three were recently exposed to single MU feedback. Experiments were carried out on 14 days in blocks of four to five consecutive days with never more than two days of break in between blocks. Each experimental session lasted approximately two hours. One subject withdrew from the experiment after only ten sessions due to time constraints. The study was approved by the ethics committee at Imperial College London (reference number: 18IC4685).

### 2.2. Data acquisition

High-density surface EMG (HDsEMG) was acquired from the tibialis anterior muscle (TA) of the dominant leg via a 64-electrode grid (5 columns and 13 rows; gold-coated; 1 mm diameter; 8 mm interelectrode distance; OT Bioelettronica, Torino, Italy). The adhesive electrode grid was placed over the muscle belly aligned to the fibre direction. In addition, EMG from the fibularis longus (FL) and the lateral and medial head of the gastrocnemius muscles (GL and GM, respectively) were recorded throughout the experiment via pairs of wet gel electrodes (20 mm interelectrode distance; Ambu Ltd, St Ives, United Kingdom) placed over the muscle belly. All EMG signals were monopolar recorded, amplified via a Quattrocento Amplifier system (OT Bioelettronica, Torino, Italy), sampled at 2048Hz, A/D converted to 16 bits, and digitally band-pass filtered (10-500Hz). The foot of the dominant leg was locked into position to allow dorsiflexion of the ankle only. The force due to ankle dorsiflexion (single degree-of-freedom) was recorded via a CCT TF-022 force transducer, amplified (OT Bioelettronica, Torino, Italy), and low-pass filtered at 33Hz. The communication between the amplifier and the computer was conducted via data packages of 256 samples (one buffer corresponds to a signal length of 125ms). All incoming EMG signals were band-pass filtered between 20-500 Hz using a 4^th^ order Butterworth filter. Bipolar derivations were extracted from the filtered EMG signals obtained from FL, GL, and GM.

### 2.3. Experimental paradigm

#### 2.3.1. Pre-experimental calibration

Subjects were instructed to perform maximum isometric dorsiflexion of the ankle to estimate the maximum voluntary contraction level (MVC). The obtained MVC was then set as a reference value for the subsequent experimental session. In a sub-MVC task, subjects were instructed to follow a 4s ramp trajectory (2.5% MVC per second) followed by a constant phase at 10% MVC of 40s. In both experiments, visual feedback of the force produced by isometric TA contractions was provided. Based on the EMG of the TA recorded during this sub-MVC task, the separation matrix used by an online decomposition algorithm was generated to extract MU discharge behaviour in real-time (see [25] for further explanation). The decomposition results were visually inspected while subjects were instructed to recruit MUs one after another based on the visual feedback provided.

#### 2.3.2. Force feedback task

After initialising the real-time decomposition algorithm, subjects were instructed to follow ramp trajectories consisting of a 10s incline (1% MVC per second) followed by a 10s plateau at 10% MVC and a 10s ramp decline (−1% MVC per second) guided by visual feedback of the force. This ramp trajectory was repeated five times with 5s rest period in between ramps. Based on the recorded force and underlying MU activity during the incline phases, the recruitment order of the decomposed MUs was estimated as suggested in [26]. The onset of MU recruitment was defined as the time when a MU started to discharge action potentials at 5 pulses-per-second (pps) or above. The averaged force values, extracted from a 100ms window centred around this onset of MU activity during all five ramps, was used to establish the MU recruitment order by ranking MUs based on their corresponding force values in ascending order. The plateau phases of the ramps were used to estimate the average DR of each MU during 10% MVC. These values were later used during the target task (see 2.3.4) to normalise the DRs. The decline phase was used to determine the force values associated with MU de-recruitment. The time point of MU de-recruitment was defined by the last action potential discharged before a MU turned “silent” for at least 1.5s. Similar to the calculation of the force level needed for MU recruitment, the force level at MU de-recruitment was estimated by the average force value extracted from a 100ms window at the offset of MU activity across all five ramps.

#### 2.3.3. MU selection

Subjects were provided with visual feedback on the ranked MU activity. In an exploration phase (approximately 10 min), subjects were instructed to recruit MU one-by-one by gradually increasing the contraction level of the TA until all identified MUs were discharging action potentials. The entire pool was divided into two sub-pools comprising the first and last recruited half of MUs, respectively. One pair of MUs with a similar recruitment threshold from each sub-pool was randomly selected (see Figure 1). Hereby, MU pairs were excluded from the selection if subjects could not recruit these two MUs one by one even after the initial exploration phase. The MU recruited first in each pair was labelled as *MU1*, while the MU recruited last as *MU2*.

**Figure 1:**
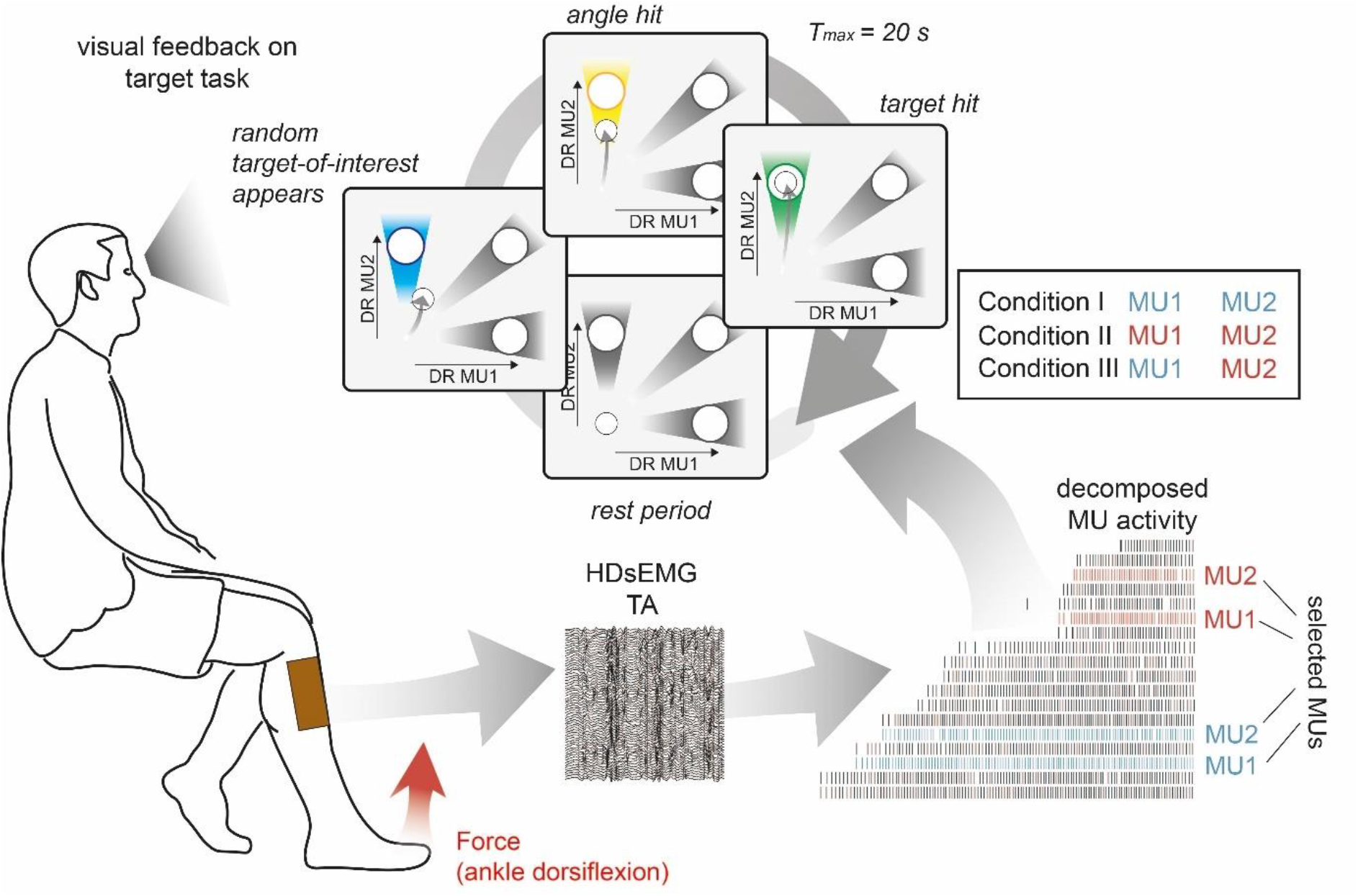
Schematic overview of the target task. HDsEMG of TA was acquired and decomposed in the underlying neural activity in real-time. Concurrently, the force due to dorsiflexion of the ankle (red arrow) and bipolar EMG of FL, GL, and GM (green arrows) were recorded. The identified MU pool was ranked accordingly to the recruitment order. Two pairs of MUs with a similar recruitment threshold were selected from the initial (blue) and the latter recruited half (red). During the target task, subjects were instructed to navigate a cursor inside a 2D space by modulating the normalised DR of MU1 and MU2. The selection of MU1 and MU2 was determined by three different conditions. In Condition I, MU1 and MU2 were coming from the low recruitment threshold pair (blue), in Condition II from the high recruitment threshold pair (red), while in Condition III, the lowest threshold MU of the low threshold pair was pooled with the highest threshold MU of the high threshold pair. During the target task, subjects were asked to stay inside the origin until the target-of-interest (blue) appeared (randomly selected). By navigating the cursor inside the angle area around the target-of-interest, subjects were granted an angle hit (yellow). The trial was terminated when either the subject managed to place and hold the cursor inside the target area (target hit, green) or more than 20s have passed since the target-of-interest appeared. In each condition, 30 targets were shown, i.e. each target ten times.

#### 2.3.4. Target task

In the main experiment, subjects navigated a cursor in a 2D space by modulating the DR of MU1 and MU2. The normalised DR of these two MUs was used to span the manifold with units ranging from 0 to 1 along both axes. As illustrated in Figure 1, this target space included three targets of equal size (radius of 10% of the normalised DR) placed along the axes (TI [1 normalised DR MU1; 0 normalised DR MU2], TIII [0; 1]) and the diagonal (TII [1; 1]). In addition, each target was framed by an angle space comprised of a triangle with one corner in the coordinate origin [0; 0] and two sides to be tangent at the circumcircle of the corresponding target (see Figure 1). Towards the coordinate origin, the angle area was cropped by a circle centred at the origin with a radius of 40% of the normalised DR. In order to navigate the cursor inside this angle space, subjects would need to generate the same discharge relationship between MU1 and MU2 as for reaching towards the target area but without matching the exact DR. For example, to place and hold the cursor inside the angle space of TI, subjects would need to keep MU1 active while MU2 inactive. However, the DR of MU1 could be different from the normalised DR of MU1, which would be required to reach the target space of TI.

The discharge behaviour of MU1 and MU2 was decomposed from the acquired HDsEMG in real-time. The obtained DRs were averaged over the preceding eight buffers (corresponds to 1s windows) and normalised by the average DR at 10% MVC of the respective MUs (see 2.3.2). The cursor movement was updated every buffer (corresponds to 125ms) and smoothed over six buffers (corresponds to 750ms) using a moving average. In total, the moving average on the DRs and the cursor position resulted in a weighted average of an effective window of 1625ms while the emphasize was on the most recently recorded second. In an initial familiarisation phase, subjects could freely move inside the target space and explore different control strategies. During this period, the gain along each axis was set manually to enable subjects to reach the target areas without overexerting themselves to prevent symptoms of muscle fatigue. On average, across all subjects, the gain was increased to 1.15 ± .01 for MU1 and 1.16 ± .01 for MU2, respectively.

After the familiarisation with the target environment, subjects were asked to rest in the coordinate origin. Once the target-of-interest appeared, indicated in blue colour, a trial started, and subjects were instructed to navigate the cursor as quickly and as directly as possible into the target area. If the cursor was kept inside the target-of-interest for at least seven consecutive buffers (corresponds to 875ms; one buffer size longer than the moving average), subjects were granted a target hit, and the trial ended. The trial was also terminated if subjects failed in navigating and holding the cursor inside the target area within 20s. If the subjects kept the cursor inside the angle area of the target-of-interest for seven consecutive buffers before the trial was terminated, the subjects were granted an angle hit. Therefore, in a single trial, subjects could achieve both an angle and target hit. A target hit was indicated via colour change of the target-of-interest to green, an angle hit to yellow, and a failure (no target nor angle hit within 20s) to black. Once the trial ended either after 20s or a target hit, the subject was instructed to navigate the cursor back into the coordinate origin and rest there for at least 2s. The entire trial cycle is illustrated in Figure 1. In total, every target was presented ten times in randomised order. Moreover, this target task was repeated three times for three different conditions. In Condition I, MU1 and MU2 were coming from the lower threshold pair while, in Condition II, they were taken from the higher threshold pair. In Condition III, MU1 was the lower threshold unit from the low-threshold pair and MU2 the higher threshold MU from the high-threshold pair (see Figure 1). After the target task was completed, subjects repeated the force task (see 2.3.2).

### 2.4. Analysis

#### 2.4.1. Force task

EMG of TA, FL, GL, and GM acquired during both force task before and after the target task were rectified and low-pass filtered at 10Hz with a 4^th^-order Butterworth filter. Similar to the estimation of the force level at the on- and offset of MU activity (see 2.3.2), the average global EMG values to MU de-/recruitment of all muscles were calculated. The average value inside a 100ms window centred around the time point of recruitment and de-recruitment for each MU and muscle was calculated across all ramps. This was separately repeated for all values acquired before and after the target task. Three subjects did not repeat the force task after the target task and only followed a single ramp at the beginning of the experiment. Moreover, no de-recruitment threshold was determined for those subjects.

#### 2.4.2. Target task

If subjects failed to navigate and hold the cursor for 875ms inside the target-of-interest within the 20s-time window, the nearest miss was calculated. The nearest miss was defined as the average cursor position over 875ms with the shortest Euclidean distance towards the centre of the target-of-interest. In addition, as previously described in [27], unintended hits were used as a metric to assess the effectiveness of subjects directly hitting the target-of-interest without unintentionally hitting unselected targets before. Therefore, an unintended hit was classified as the case when subjects navigated and held the cursor inside an unselected target for at least 875ms. Unintended hits of the same target could occur multiple times within a single trial if the cursor re-entered the unselected target on several occasions before the trial was terminated. Similarly, unintended angle hits were counted when the cursor was navigated into unselected angle areas, respectively.

TI and TIII required the sole activation of either MU1 or MU2. To assess subjects’ performance in navigating the cursor towards these two targets even when neither the target nor the angle area was reached, a new performance metric was introduced. This performance metric was defined as:

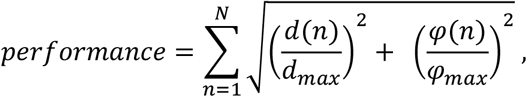

where *d*(*n*) is the Euclidean distance between the centres of the cursor and target-of-interest and *φ*(*n*) the angle between the cursor and the target-of-interest at the buffer *n. N* is the total number of buffers recorded in one trial. For TI and TIII, *d_max_* was set as the Euclidean distance between the centres of TI and TIII, and *φ_max_* to 90°. For TII, the target along the diagonal, *d_max_* was set to the distance between the centre of TII and the origin, and *φ_max_* to 45°. When the cursor was held in the origin, i.e. no activation of either MU1 or MU2, *φ*(*n*) was set to *φ_max_*. By summing across all recorded buffers, this metric incorporates the time-to-target and favours those trials in which the cursor was kept close to the target-of-interest even when neither the angle nor the target area was hit. To scale the *performance values* between 0 and 1, the obtained result was normalised with the *worst* and *best performance values* estimated per target. The *best performance value* for each target across subjects was obtained by simulating cursor movement based on artificially generated MU discharge patterns that match the required activation of MU1 and MU2 to hit the respective target. For example, the *best performance* for TI was estimated based on discharge behaviour for MU1 that matched a normalised DR equal to 1 and MU2 equals 0. The performance value during the idealised cursor movement until the target hit was used as the corresponding *best performance value*. For the *worst performance value*, the performance was calculated as if the cursor was kept in the origin for the entire 20s.

The described metrics were calculated across all conditions and subjects. Three subjects started with Condition III only from day 10 onwards.

### 2.5. Questionnaire

After every condition, subjects were provided with a questionnaire. Subjects were asked to indicate the level of control they had over MU1 when reaching towards TI, over MU2 when going towards TIII, and both MUs when reaching towards TII. Moreover, it was assessed whether subjects felt using a concrete strategy when going to the selected target-of-interest and how cognitively demanding it was to control the MUs together and independently. When applicable, they were asked to explain their strategy. Three subjects did not fill out the questionnaire.

### 2.6. Statistics

Statistical analysis was conducted using SPSS (IBM, Armonk, NY, USA) and Matlab (Version 2018b, The Mathworks, Inc., Natick, MA, USA) for the linear mixed model analysis. The threshold for statistical significance was set to p < .05. To avoid accumulation of Type I errors, non-parametric tests were used to assess the relationship between variables [28]. To compare recruitment and de-recruitment thresholds, a linear mixed model with the difference between recruitment and de-recruitment threshold as dependent variable, a fixed effect intercept and a participant specific random intercept was applied using restricted maximum likelihood estimation. Significance of the fixed effect was assessed by an F-test using Satterthwhaite’s approximation for the degrees-of-freedom. For analysing the improvement in target and angle hit rate, performance across and for each target, as well as the relationship between the performance of reaching each target and the difference in recruitment threshold between MU2 and MU1, two-sided Wilcoxon signed-rank tests were used. Comparison of mean characteristic forces during indirect hits of TIII were conducted via Friedman test. A two-sided Wilcoxon signed-rank test was used for post-hoc analysis.

## 3. Results

On average, 11.04 ± 3.34 MUs were reliably decomposed per subject. An identified MU pool from a subject sorted based on recruitment order is shown in Figure 2A. As indicated by the green and red marks, the order in which MUs are de-recruited often differed from the recruitment order. For example, once recruited, a MU could keep discharging action potentials even when the exerted force level was below the initial recruitment threshold. For the target task (see 2.3.4), two pairs of MUs with a small difference in recruitment threshold were selected out of the entire pool. Each pair was selected either from the first or from the last recruited half of the MU pool. Figure 2B visualises the recruitment thresholds of these selected MUs. Within pairs, MU1 was recruited before MU2 and the lower threshold pair (blue) was recruited before the higher threshold one (red). Recruitment and de-recruitment threshold of the selected MUs showed a strong relationship across days and subjects (Spearman correlation coefficient, R = .62, p < .001; see Figure 2C). On average, the de-recruitment threshold was −.83 ± 2.08 % MVC smaller than the recruitment one, and in 64.73% a selective MU was de-recruited at a force level below its initial recruitment threshold. The overall observed effect, however, was weak (see 2.6, p = .21).

**Figure 2:**
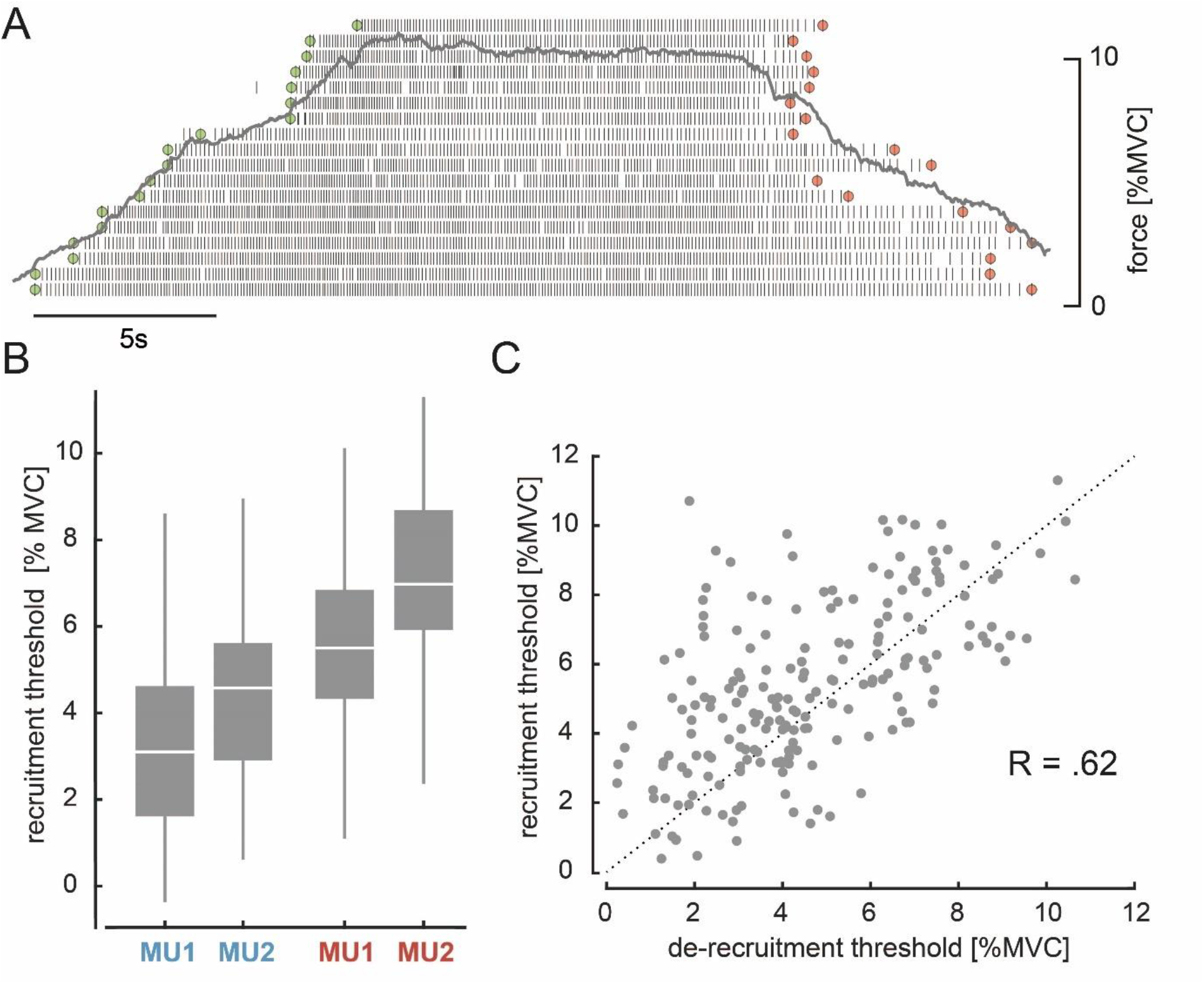
MU recruitment and de-recruitment. A: The identified MU pool ranked on the recruitment order of one representative subject is shown with the underlying force profile (grey). Time points of recruitment (green) and de-recruitment (red) for each MU are marked. B: Recruitment level for MU1 and MU2 of the lower (blue) and higher threshold pair (red) across all subjects and days are shown with their median and quartiles. C: Recruitment and de-recruitment threshold for the selected MUs across all days showed a significant relationship (p < .001). Dashed line indicates the diagonal. The three subjects for whom no de-recruitment thresholds were determined were neglected in this correlation analysis.

During the target task, subjects were asked to navigate a cursor inside a 2D space by modulating the DR of MU1 and MU2. Three different targets inside the 2D plane were used to encourage subjects to activate both MUs independently despite their different position within the recruitment order. For example, to reach TIII, subjects must keep the higher threshold unit MU2 active while keeping the lower threshold unit MU1 off. Figure 3A shows, as an example, the average cursor position during target hits and nearest misses for each target-of-interest across conditions towards the beginning and end of training for a single subject. At the beginning of the task, the subject failed in the majority of trials to place and hold the cursor inside the target-of-interest. With training, the ability to place the cursor inside the designated target area improved. As shown in Figure 3A, TII was hit in all 30 trials, and in only four trials, the subject could not hit TI. Moreover, the nearest misses for TIII in the twelfth training session were closer to the target centre than on day 1. This improvement in hitting targets and angles over several training days was observed across all subjects (see Figure 3B). The target hit rate improved from the first to the last day of training from 41.19 ± 17.76% to 67.04 ± 18.17% (two-sided Wilcoxon signed-rank test, p = .028). A similar trend was observed for the angle hits with an improvement from 64.37 ± 13.15% to 81.11 ± 9.84% (two-sided Wilcoxon signed rank test, p = .016). According to the distanced-based performance metric (defined in 2.3.4), the performance per subject across targets and conditions improved from the first to the last day of training from .56 ± .09 to .69 ± .10 (two-sided Wilcoxon signed rank test, p = .016). Despite this clear improvement in performance across days, the ability to move the cursor towards the target-of-interest enhanced differently across targets (see Figure 3C). The performance in hitting TI did not significantly improve from the first day of training to the last day (from .67 ± .18 to .68 ± .11; two-sided Wilcoxon singed rank test, p = .938). Similarly, no significant improvement was observed for TII (from .69 ± .11 to .80 ± .13; two-sided Wilcoxon signed rank test, p = .109). However, a significant improvement in performance was detected when subjects were asked to move towards TIII (from .33 ± .12 to .59 ± .17; two-sided Wilcoxon signed rank test, p = .016)

**Figure 3:**
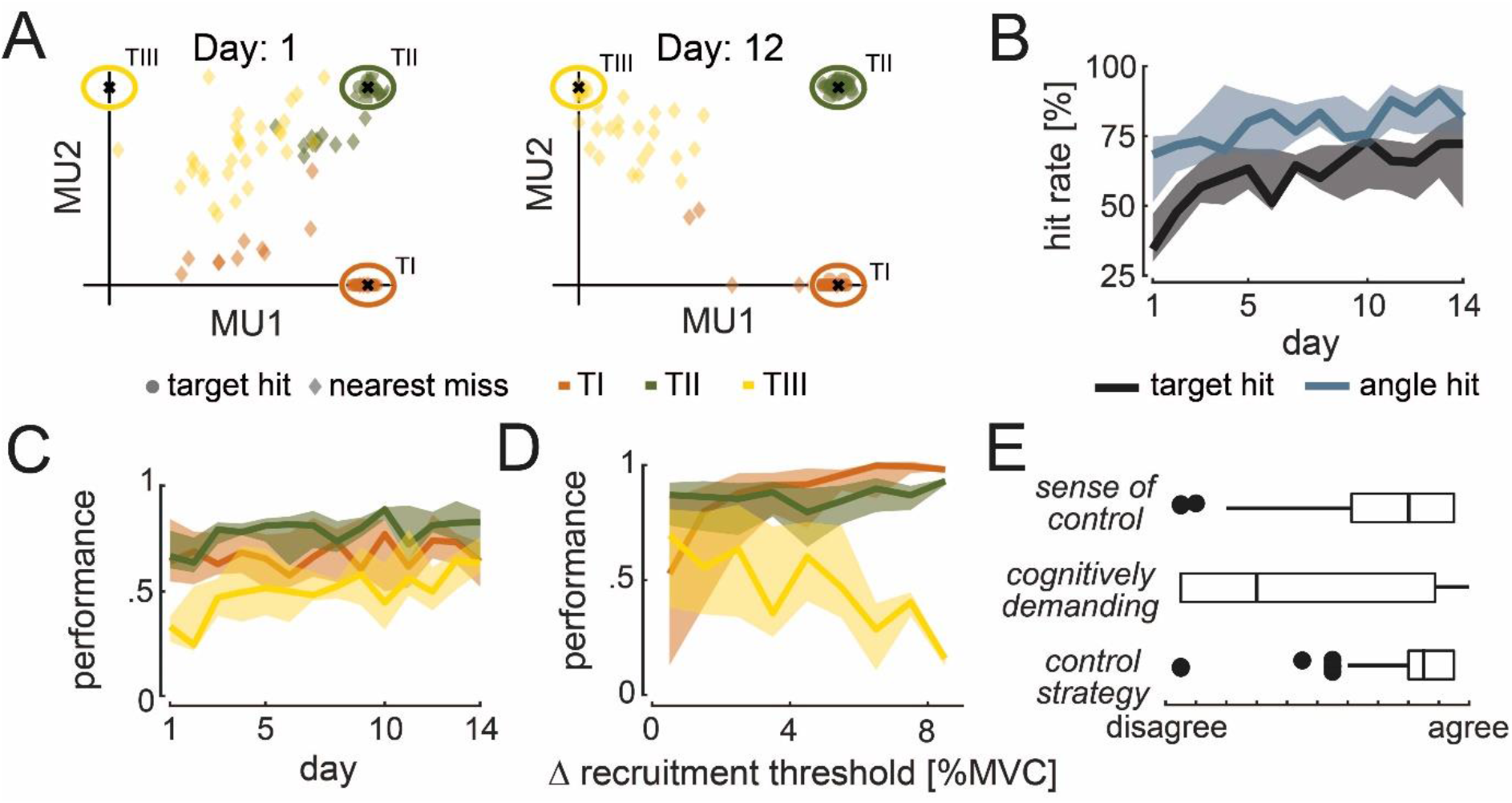
Cursor movement and performance during target task. A: Average cursor position during target hits (circle) and nearest misses (diamond) across conditions are shown for the first and twelfth day of experiments for TI (orange), TII (green), and TIII (yellow) of one subject. B: Target (black) and angle hit rate (blue) across subjects, conditions and targets-of-interest are shown with their medians (solid line) and 25% and 75% quartiles (shaded areas) across days. C: Performance values across subjects and conditions for TI (orange), TII (green), and TIII (yellow) are shown with their medians (solid line) and 25% and 75% quartiles (shaded areas) across days. D: Performance values corresponding to the difference in recruitment threshold between MU2 and MU1 are shown across subjects and conditions for the last five days of training. The median (solid line) and 25% and 75% quartiles (shaded area) for TI (orange), TII (green), and TIII (yellow) are illustrated in steps of 1% MVC. E: Subjective experience of controlling MU2 when reaching towards TIII based on the questionnaire response during the last five days of training across subjects and conditions are shown by their medians and quartiles.

Taken together, target and angle hits, as well as the performance metric, indicate that subjects improved in navigating the cursor towards the target-of-interest across days. However, the main improvement was observed for reaching TIII. Moreover, subjects experienced a steep learning curve at the beginning of the experiment, while the learning rate seemed to decrease towards the end. For this reason, further analysis only focuses on the last five days of training to avoid additionally induced variability by greater learning rates at the beginning of training.

The performance of reaching each target did not solely depend on the target’s position but also on the difference in recruitment threshold within the selected MU pair, i.e. the force difference between the onset of activity between MU2 and MU1 measured during the initial force task (see 2.3.2). Figure 3D illustrates the performance over the within-pair difference in recruitment threshold for each target-of-interest. Performance per subject in reaching TI significantly increased from selected MU pairs with a small difference in recruitment threshold (0-1% MVC) to those with a high difference (3-10% MVC) from .56 ± .19 to .87 ± .10 (two-sided Wilcoxon signed rank test, p = .016). To reach TI, only MU1 must be active. Therefore, this result indicates that subjects performed better in keeping only MU1 active while not activating MU2 when the difference in their recruitment threshold was high. This may be due to less accuracy needed in the force generated when the recruitment threshold difference is large. For example, if MU1 gets recruited at 2% MVC while MU2 at 5% MVC, the subject could potentially exert any force between 2 and 5% MVC to keep only MU1 active without MU2 to ultimately hit TI. A more precise force level needs to be generated when this difference is smaller. For TII, no dependency in performance and the within-pair difference in recruitment threshold was detected (from .81 ± .12 to .79 ± .12, two-sided Wilcoxon signed rank test, p = 1). On the contrary, the performance in reaching TIII decreased significantly for larger differences in recruitment threshold within the selected MU pair from .66 ± .15 to .45 ± .13 (two-sided Wilcoxon signed rank test, p = .016). This indicates that subjects experienced difficulties in keeping MU2 active while MU1 is inactive in order to reach TIII when their difference in recruitment threshold was large. During the last five days of training, the majority subjects reported that they felt having control over MU2 when reaching towards TIII (14% of cases subjects indicated having no control over MU2; see Figure 3E). Moreover, they declared the usage of a clear strategy to establish such control, which varied in cognitive demand across subjects and days. In 95% of all cases, this control strategy was described as a rapid increase in force due to dorsiflexion of the ankle followed by a slow release until the cursor moves towards the vertical axis.

All subjects improved their performance during training. To better understand which strategies have emerged, ultimately enabling subjects to recruit and de-recruit single MUs, we analysed the cursor trajectories during the task. The cursor movement for one subject during the last day of training (Condition I) for all 30 trials is visualised in Figure 4. While the subject was able to hit all targets before the trials ended, the cursor trajectories did not always mimic the straight path, to the target centre. When asked to move towards TI, the subject moved the cursor along the horizontal axis. In trials 1, 2, 3, 5, 9, and 10, the subject could not hit TI in the first attempt but returned to the origin to then move towards the target centre directly. For TII, in all cases, the subject moved directly along the diagonal towards the target-of-interest. For TIII, however, instead of moving directly towards the target centre, the subject moved the cursor towards TII first and then towards the vertical axis to finally hit TIII. This observation is in line with the descriptions provided by the questionnaire (see Figure 3E), i.e. increase in force to activate both MUs, followed by a decrease in the force until MU1 switches off, and ultimately adjusting the force level to move along the vertical axis towards the target centre. By analysing the unintended target and angle hits (see Figure 5A), the probability that the cursor was moved towards unselected targets while trying to hit the target-of-interest was quantified.

**Figure 4:**
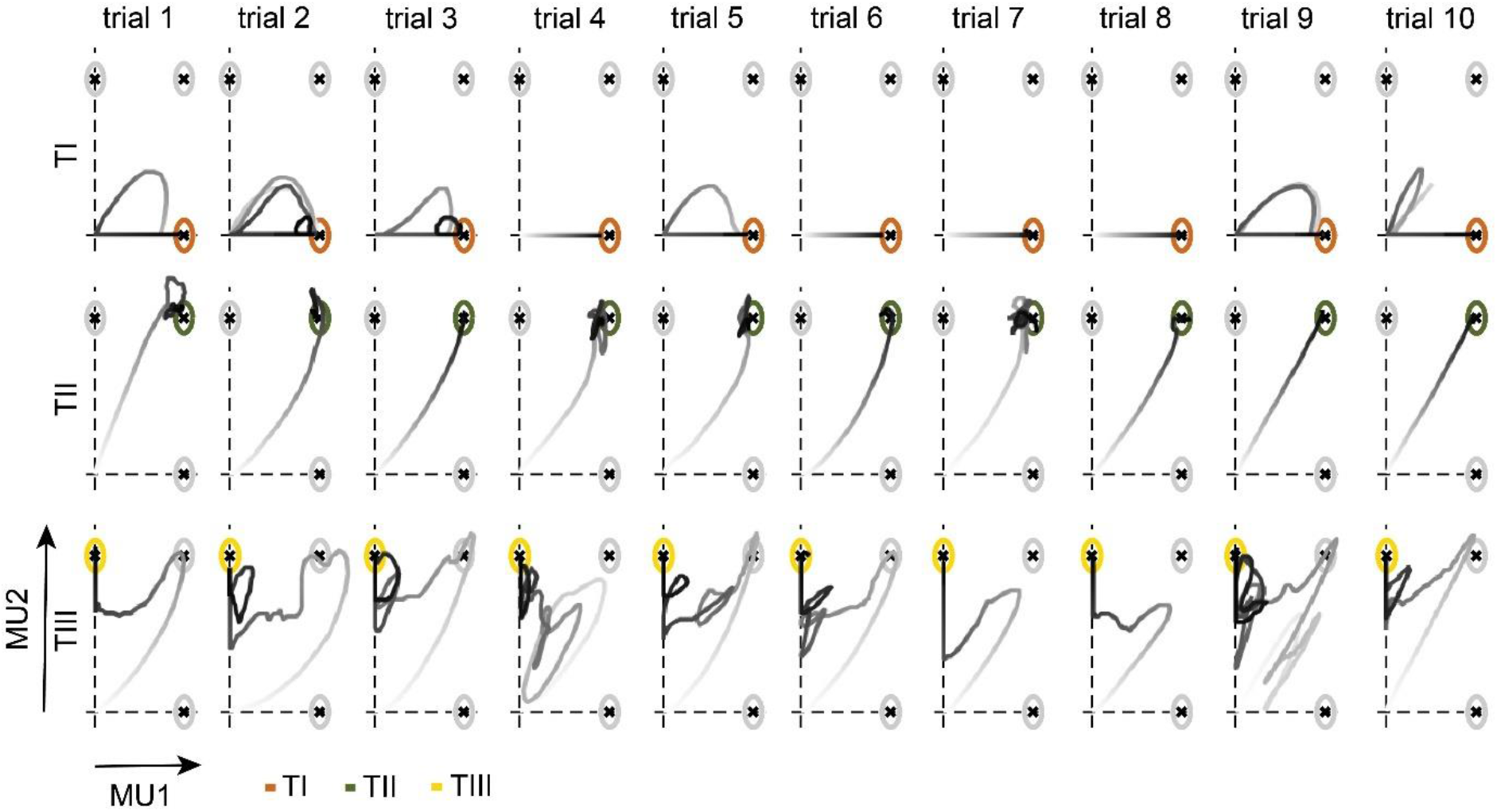
Cursor trajectories of one subject during the target task. Cursor movement towards TI (orange, top), TII (green, centre), and TIII (yellow, bottom) in each trial for Condition I on the 14^th^ day of one representative subject is shown. Trial 1 to trial 10 indicate the first to the tenth appearance of each target-of-interest. The grey intensity of the cursor trajectories increases over time within the trial.

**Figure 5:**
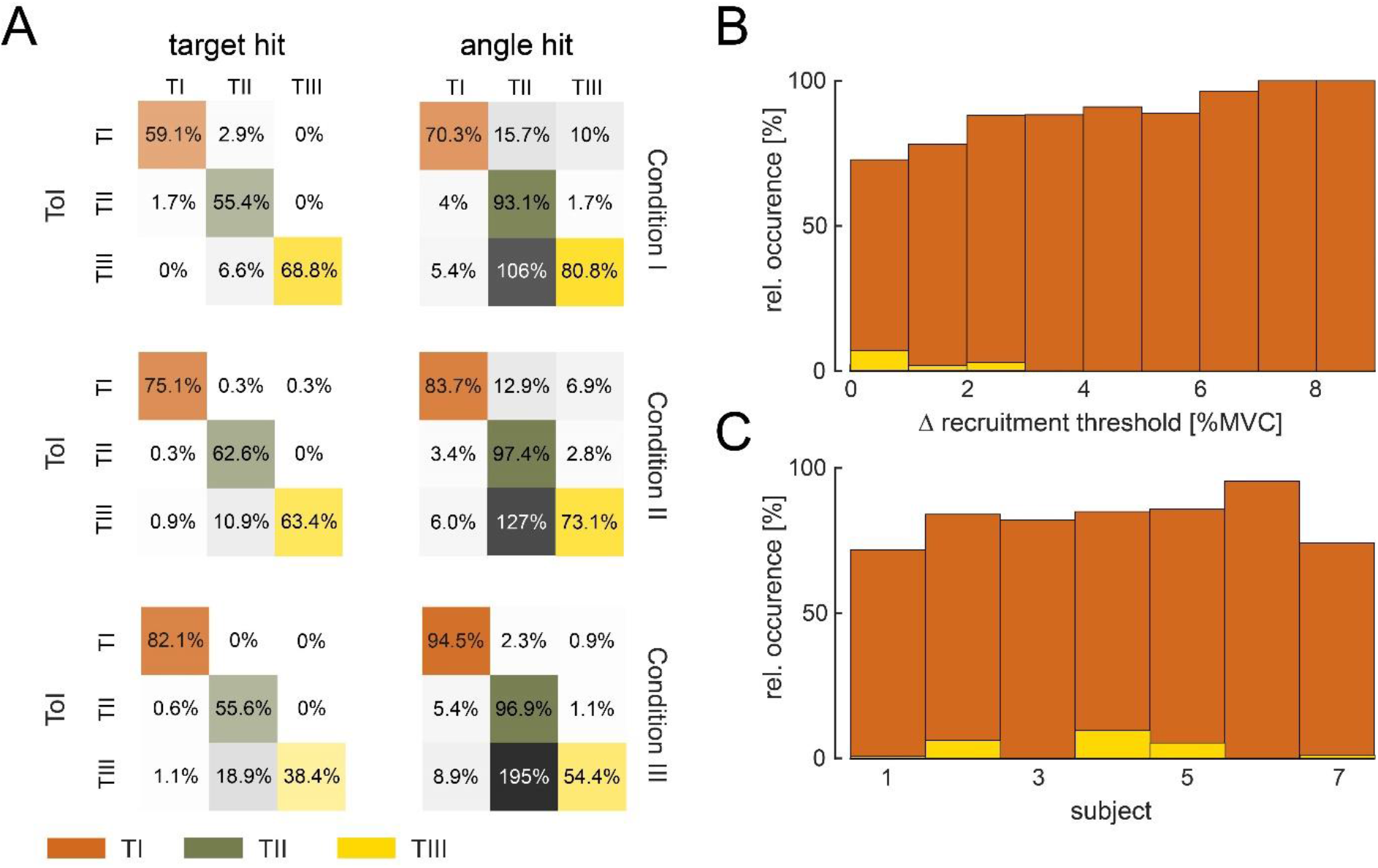
Movement towards targets-of-interest. A: Relative hit rate of intended and unintended hits (grey) of targets and angles are shown for the targets-of-interest (ToI) TI (orange), TII (green), and TIII (yellow) during the last five days of training across subjects for Condition I (top), Condition II (centre) and Condition III (bottom). Please note that hit rates above 100% can be reached for unintended hits when subjects re-entered the target before the trial ended. Colour intensity corresponds to the hit rate. Relative occurrence of direct movement towards TI (orange), i.e. only activating MU1 without MU2, and TIII (yellow), i.e. only activating MU2 without MU1, during successful attempts (at least angle hits), are shown with respect to the difference in recruitment threshold between MU2 and MU1 (B) and across subjects (C).

While only a few unintended hits occurred across targets and conditions, unintended angle hits of TII happened multiple times when subjects tried to reach for TIII. In fact, across all conditions, almost all subjects conducted most of their unintended hits when aiming for TIII (with unintended hits in TII). Only one subjects had a higher rate of unintended hits when aiming for TII (with unintended hits in TI) in Condition I. Therefore, the strategy to reach TIII, i.e. moving towards TII first, as illustrated in Figure 4 and interpreted by the questionnaire answers, can be observed across subjects. Moreover, the few unintended hits when reaching towards TI and TII suggest that subjects established control strategies that allowed for a direct movement towards the target-of-interest. These clear control strategies, as well as the difference in learning rate across targets, suggest that subjects were able to activate MU1 alone (to reach TI), MU1 and MU2 together (to reach TII) but could not volitionally activate MU2 before MU1 (TIII). In fact, during the last five days of training, 76%, 78%, and 94% of all successful attempts, i.e. at least an angle hit, of going towards TI were achieved without activating MU2 once during the trial while it was only 7%, 2%, and 1% for TIII (MU2 only without MU1) in Condition I, II, and III, respectively. The percentage of these direct movements towards TI and TIII with respect to the difference in recruitment threshold within the selected pair and subjects is shown in Figure 5B. While direct movements towards TI increased with a larger difference in recruitment threshold, direct movements towards TIII were very rare and only possible with MUs recruited at very similar force levels, i.e. with small differences in recruitment thresholds that led to variable recruitment orders given sudden excitatory inputs at the beginning of the trials. Also, all subjects moved directly towards TI in more than 70% of all successful attempts. Although, only three subjects navigated the cursor directly towards TIII in more than 5% of all successful attempts (see Figure 5C).

In these rare cases in which direct movements towards TIII ended, at least, in an angle hit, the level of force and global EMG of TA, FL, GL, and GM at the time point of recruitment of MU2 was compared with the corresponding values obtained at normal recruitment of MU2 during the initial force ramps (see 2.4.1). During direct movement towards TIII, MU2 was recruited on average at a 52.94 ± 27.72% lower force level than during ramp recruitment. Also, the global EMG values at recruitment of MU2 were, on average, slightly lower during direct movements towards TIII (TA: −2.46 ± 16.53%; FL: −8.83 ± 17.74%; GL: −8.98 ± 19.97%; GM: −6.29 ± 19.98%).

However, in the vast majority of cases, target hits of TIII were not achieved by direct movements towards the target centre. Instead, subjects used a three-stage approach to place the cursor inside TIII, as observed in Figure 4 (for example, TIII, trial 7). First, subjects navigated the cursor along the diagonal towards TII (stage one) before in the second stage moving towards the vertical axis. In the third stage, subjects manoeuvred the cursor along the vertical axis inside TIII. The discharge rate of MU1 and MU2 and the exerted force during indirect movement towards TIII are shown for a representative subject in Figure 6A. During the first stage, the subject increased the force to orderly recruit MU1 and MU2. Once both MUs were active, the subject decreased the force level in stage two to a minimum so that MU1 stopped firing while keeping MU2 active. In the third stage, the force was slightly increased to match the necessary DR of MU2 to reach TIII without re-activating MU1. To assess whether such force modulation during indirect hits of TIII could be observed across subjects and conditions, characteristic forces (due to ankle dorsiflexion) for each stage were compared in Figure 6B. The characteristic force during stage one was the mean force in a 100ms window around the maximum force when both MUs were active, i.e. DR greater than 5pps. In stage two, the characteristic force was estimated by averaging the force inside a 100ms window at the minimum force level after switching off MU1 while MU2 continued firing action potentials. In stage three, the characteristic force was set as the mean force during the hold period preceding a target hit of TIII. Across conditions, all subjects used significantly different force levels during each stage (Friedman, p < .001, two-sided Wilcoxon signed rank test Bonferroni corrected, always p < .05). During stage one, the exerted force level was the greatest while being reduced to a minimum in stage two, before being slightly increased again in stage three. Furthermore, for the four subjects for whom the de-recruitment threshold was determined, 24.32% of all indirect target hits of TIII were achieved while MU2 was de-recruited before MU1 during the initial force ramps. This indicates that this three-stage approach also worked for pairs of MUs for which the de-recruitment threshold was not reversed to the recruitment one.

**Figure 6:**
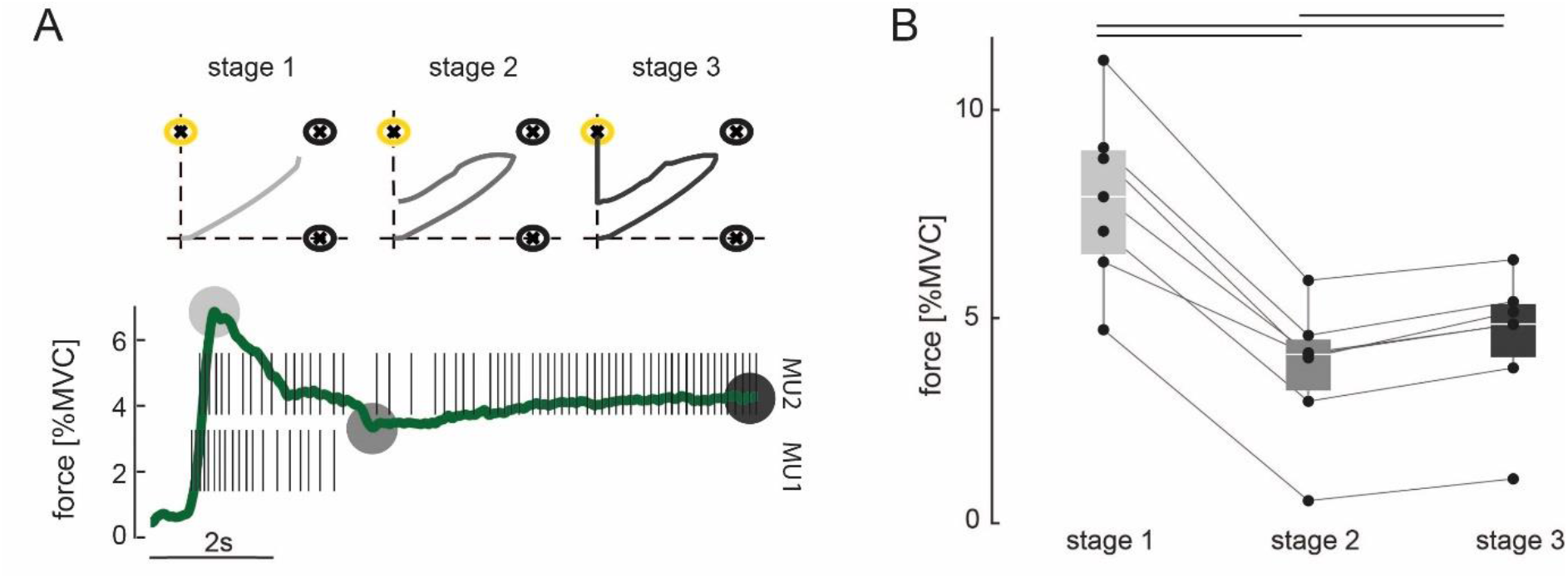
Three-stage approach to hit TIII. A: Force due to ankle dorsiflexion (green) and the discharge behaviour of the selected MU pair during a successful attempt of hitting TIII (yellow) for a representative subject. The subject used a three-stage approach to achieve the target task: stage 1: increasing the force to orderly recruit both MUs; stage 2: reducing the force until MU1 stops firing while the cursor is placed along the vertical axis; stage 3: slightly increasing the force again to manoeuvre the cursor inside TIII. Grey circles mark the characteristic force values of each stage. Stage 1: maximum force while both MUs are active; stage 2: minimum force after MU1 stopped firing; stage 3: force during hit of TIII. The corresponding cursor movement for each stage is shown on the top. Grey intensity increases with stages. B: Characteristic forces (due to ankle dorsiflexion) are shown with their median and quartiles at each stage of control for all subjects across all conditions during all TIII hits in the last five days of training. Each dot represents a subject, and corresponding values are connected via the lines. Black bars indicate a significant difference with p < .05.

The force, global TA, FL, GL, and GM EMG values in 100ms window centred around the onset of MU activity were compared before and after the experiment to investigate potential changes in the recruitment order due to single MU modulation (see 2.4.2). A subtle decrease in force (−.08 ± 2.33 % MVC) and global TA EMG (−0.74 ± 15.94%) were identified. However, the global EMG from the lower leg muscles not used for the MU decomposition increased slightly (FL: 7.65 ± 27.64%; GL: 16.52 ± 37.26%; GM: 18.54 ± 24.68%). These changes indicate that the overall recruitment order did not change critically due to the imposed single MU modulation. The increase in activity in the lower leg muscles not directly involved in ankle dorsiflexion relative to the agonist muscle might be explained by induced fatigue towards the end of the experiment [29].

## 4. Discussion

Volitional and flexible control of single MUs could revolutionise neural-interface applications. Here we used real-time biofeedback on single MU activity to encourage subjects to learn independent control of pairs of MUs. Our results showed that subjects could gain control over four MUs from a single muscle. The control strategies that emerged, allowing for selective MU control, were limited by the presence of a common input to the MU pool. Therefore, subjects did not exploit potential neural structures with selective inputs to individual MUs.

In this study, the identified MU pools were ranked based on their recruitment order. We have shown that the de-recruitment threshold does not always match the recruitment one. This might be due to intrinsic neuro-modulatory mechanisms triggered following the activation of a MU which can disrupt the simple dependency of recruitment order from neuron size and input received [4]. One potential mechanism contributing to this neuromodulation in motor neurons are persistent inward currents (PICs) [30]. PICs can alter the excitability of motor neurons which may lead to a self-sustained firing of action potentials and thus to de-recruitment at a force level lower than the recruitment threshold, i.e. recruitment de-recruitment hysteresis [30]. In this study, we examined the behaviour of pairs of MUs at very low forces, i.e. less than 10% MVC. These low-threshold MUs active at such force levels may be more prone to the effect of sustained PICs [31], which might explain the hysteresis between recruitment and de-recruitment threshold observed for most MUs examined in this experiment. Moreover, the effect of PICs is pronounced when a MU is repeatedly recruited and de-recruited [32]. The amount of unintended angle hits of TII preceding an angle hit of TIII suggested that subjects required multiple attempts to switch off MU1 while keeping MU2 active. Hence, MU2 was recruited and de-recruited repeatedly within a short amount of time. Although, we did not measure PICs, these repeated activations seemed to increase the excitability of MU2, ultimately, resulting in sustained firing of MU2 while MU1 was deactivated. Furthermore, the low rate of unintended hits in TII or TIII (both require the activation of MU2) indicated that this effect on the excitability of MU2 was diminished by the short breaks in between trials, which is in agreement with previous observations on PICs [32].

An inhibitory input is needed to extinguish the impact of PICs on the MU discharge behaviour. Such a continuous background inhibitory signal leads to the reversal of MU activity to the initial state once a MU stopped firing for a prolonged amount of time. This explains why MU recruitment was not critically altered even after extensive single MU control during this experiment. The very rare cases in which an activation of the higher threshold MU before the lower threshold one occurred (direct movement towards TIII) and thus recruitment of MU2 at a lower force as during the initial ramp contractions may be explained by an incomplete extinction of the PIC effect.

During a progressive increase in force, recruitment depends only on the MU anatomy and the input received. If humans can learn to leverage potential structures in the CNS that allow selective inputs to MUs [24], changes in the recruitment order during this initial phase should be expected. However, it is important to underline that a conclusion of flexible control based on changes in MU recruitment cannot be drawn for time intervals that follow an activation of the MU. In these cases, the de-recruitment of a MU at a force level different from the recruitment threshold could be incorrectly interpreted as an alteration of the recruitment order. Presumably, such changes result from the relative intrinsic excitability of the motor neurons which override the sole impact of the received synaptic input on the recruitment order. Therefore, a direct proof of altered MU recruitment as a consequence of independent input to different MUs needs to be provided during the initial activation phase, i.e. a MU with higher recruitment threshold activated before a MU with lower threshold without preceding activations. It is also worth mentioning that this proof should further include MUs with sufficiently different recruitment thresholds since synaptic noise may influence the relative recruitment order for MUs of very similar thresholds [4].

We did not get results supporting a *general* flexible control of MUs, i.e. volitional activation of higher threshold MUs before lower threshold ones at initial recruitment. However, flexible control of individual MUs could still be a framework explaining how subjects were able to reach the different targets in the 2D space. If this was the case then, since control would be achieved only after MUs were recruited, this would imply that flexible control is state-dependent: it can only be achieved in the context of previously contracted muscle fibres. Such state-dependency restricts the possible neural strategies that could allow flexible control of MUs. One possible strategy consistent with such state-dependent control of individual MUs could be relying on an input signal to MUs not directly linked to motor function and non-homogeneously distributed among the MU pool. For example, cortical oscillations could meet these criteria if descending projections to large and small MUs in a pool differ [27], [33]. Such kind of inputs to MUs could provide a certain degree of flexibility to volitionally control subgroups of MUs in a muscle. Future studies are needed to test this hypothesis.

Throughout the 14 days of training, subjects were asked to modulate the DR of MU pairs independently to navigate a cursor as quickly as possible into different targets inside a 2D space. The target and angle hit rates indicate that subjects could achieve control over these single MUs. Moreover, the angle hit analysis in Figure 5A, for example, revealed that during the last five days of training, subjects were able to produce the necessary activation pattern in the majority of trials despite differences in initial and within-pair recruitment threshold of the selected MU pairs. Although subjects repeated the target-tracking task with different sets of MUs every day, they consistently reported the use of the same control strategies across days. Hence, these findings suggest that subjects learned to establish universal control strategies that allowed for the achievement of the target task for various combinations of MUs. To reach TI or TII, subjects used precise force control to exert either a low enough force that only the low threshold MU MU1 turned active (TI) or a force above the recruitment threshold of both MU1 and MU2 (TII). These were natural tasks that corresponded to a physiological activation of the two MUs. When asked to move towards TIII, subjects mainly mimicked the trajectory of TII first, i.e. activating both MUs, followed by the second stage of control in which the force level was reduced until the lower threshold MU turned off by leveraging the mismatch in the de-recruitment thresholds. In order to then place and hold the cursor inside TIII, subjects gradually increased the force again without re-activating MU1. This second and third stage of control were possible in principle by maintaining a common ionotropic input to the MU pair combined with neuromodulatory input, as described above, even when MU2 was initially de-recruited before MU1. If a direct activation of MU2 would have been possible as it was for MU1, subjects would have chosen to mimic a cursor trajectory along the direct path from the origin to the target centre, as observed for both TI and TII (see Figure 4). This almost never occurred in the hundreds of trials tested. Therefore, the sole activation of a higher threshold MU was only possible by exploiting the history-dependent activation of MUs, i.e. exclusive firing of MU2 follows the combined activation of MU1 and MU2. The results suggest that this three-stage approach to achieve a hit in TIII as quickly as possible was feasible for the subjects while a more efficient strategy of directly activating MU2 without a preceding activation of MU1 was not.

It has been previously shown that subjects can learn to control MUs independently when exposed to biofeedback on the discharge behaviour [2], [23]. However, in these previous investigations, the subjects were allowed for movements along multiple directions. For example, Formento et al. studied flexible control of MUs of the biceps brachii muscle while subjects were allowed to perform movements along two directions, i.e. elbow flexion and forearm supination [2]. Such variations in force directions [34], [35] but also other motor behavioural changes, including alternations in postures [36], and contraction speed [15], are well known factors that impact the recruitment order. Similar changes in a MU pool’s discharge activity imposed by such behavioural changes were recently confirmed in non-human primates [24]. Therefore, the reported observations may be triggered by small compensatory movements rather than being the result of a dedicated and volitionally controllable synaptic input to individual MUs. Indeed, independent control of individual MUs would imply that a MU can be controlled independently *of all other MUs*. The fact that a pair of MUs can be controlled independently when varying the task does not imply that the two MUs are independently controlled in absolute terms. They are simply independently controlled with respect to each other. For example, in some tasks or in some conditions, they may be part of different groups of MUs receiving two different common inputs [10].

A recent study in humans provided evidence for the existence of MU pool synergies similar to the functional grouping of muscles involved in a single movement [10]. The CNS may send a common input to these MU pool synergies, which are not per se limited to innervating only a single muscle [37]. In our experiment, we chose a simple case of a MU pool constituting a functional group during ankle dorsiflexion, i.e. MUs innervating the TA. During more complex tasks, for example, movement along multiple directions, the CNS would send different common inputs to certain numbers of groups of MUs. While the synergistic organisation of MUs might be flexible across tasks, e.g. movement along multiple directions, it remains yet to be explored if the input to a single functional MU group can be changed volitionally from common to individual while the performed task is maintained. Hence, it is crucial that initial conditions during MU recruitment, such as posture, contraction speed, and force direction, are kept constant when flexible MU recruitment is investigated. MU activation based on changes in behaviour, or the performed motor task, does not indicate that subjects can volitionally trigger MU activity by a selective synaptic input as it is possible for their cortical counterparts. Furthermore, these constraints need to be considered in neural-machine interface applications relying on flexible MU control. Possible extracted control signals may depend on behavioural changes and not on a designated descending control command and, therefore, may not be valid anymore if aimed to restore motor function when these behavioural changes cannot be triggered by the subject. Similar constraints effectively apply for augmentation when the aim is to extend the degrees-of-freedom that a human can volitional control, i.e. adding supernumerary degrees-of-freedom to the natural ones [5], [6]. In such cases, if the control of a supernumerary degree-of-freedom is based on single MU activation, this activation must be uncoupled from motor behaviour to ensure coordination between natural and supernumerary effectors. This would correspond to breaking common input into multiple inputs. Nevertheless, even based on behavioural changes, single MU control can be a resource for specific neural-interface applications, for example, in the absence of any additional motor information or to augment a specific motor task (see [5] for task augmentation).

To summarize, we have demonstrated the ability to control up to four MUs from a single muscle using real-time feedback on single MU discharge behaviour. Furthermore, we have shown by operant conditioning that subjects learn concrete control strategies to recruit and de-recruit several MUs volitionally. These strategies exploit orderly recruitment in agreement with the *Henneman’s size principle* and a common input to their motor neurons. Conversely, the observed strategies do not leverage potential pathways that may provide selective inputs to single MUs. It is concluded that converting common input to a (synergistic) pool of motor neurons into independent input to single MUs within the same task seems extremely challenging for the CNS.

## Acknowledgement

This study was supported by the EPSRC Centre for Doctoral Training in Neurotechnology and Health and the European Commission grants H2020 NIMA (FETOPEN 899626) and H2020 TRIMANUAL (MSCA 843408).

## Author contribution

MB, JI, DYB, JE, EB, CM, and DF conceived the study. MB carried out the experiments and conducted the analysis. MB, JI, DYB, JE, EB, CM, and DF interpreted the data, MB wrote and JI, DYB, JE, EB, CM, and DF edited the manuscript.

## Declaration of interests

DF and DYB are inventors in a patent (Neural 690 Interface. UK Patent application no. GB1813762.0. August 23, 2018) and DF, DYB, JI, and MB are inventors in a patent application (Neural interface. UK Patent application no. GB2014671.8. September 17, 2020) related to the methods and applications of this work.

## References

[1] A. Holobar and D. Farina, “Noninvasive Neural Interfacing With Wearable Muscle Sensors: Combining Convolutive Blind Source Separation Methods and Deep Learning Techniques for Neural Decoding,”IEEE Signal Process. Mag., vol. 38, no. 4, pp. 103–118, Jul. 2021.

[2] E. Formento, P. Botros, and J. M. Carmena, “A non-invasive brain-machine interface via independent control of individual motor units,”bioRxiv, p. 2021.03.22.436518, Mar. 2021.

[3] D. Farina and A. Holobar, “Human-machine interfacing by decoding the surface electromyogram,”IEEE Signal Process. Mag., vol. 32, no. 1, pp. 115–120, 2015.

[4] C. J. Heckman and R. M. Enoka, “Motor Unit,” in Comprehensive Physiology, John Wiley & Sons, Inc., 2012.

[5] J. Eden et al., “Human movement augmentation and how to make it a reality,”arXiv Prepr.,Jun. 2021.

[6] G. Dominijanni et al., “Enhancing human bodies with extra robotic arms and fingers: The Neural Resource Allocation Problem,”arXiv Prepr., Mar. 2021.

[7] D. Farina, F. Negro, S. Muceli, and R. M. Enoka, “Principles of Motor Unit Physiology Evolve With Advances in Technology,”Physiology, vol. 31, no. 2, pp. 83–94, 2016.

[8] D. Farina, F. Negro, and J. L. Dideriksen, “The effective neural drive to muscles is the common synaptic input to motor neurons,”J. Physiol., vol. 592, pp. 3427–3441, 2014.

[9] F. Negro, U. Ş. Yavuz, and D. Farina, “The human motor neuron pools receive a dominant slow-varying common synaptic input,”J. Physiol., vol. 594, no. 19, pp. 5491–5505, 2016.

[10] S. Tanzarella, S. Muceli, M. Santello, and D. Farina, “Synergistic Organization of Neural Inputs from Spinal Motor Neurons to Extrinsic and Intrinsic Hand Muscles,”J. Neurosci., p. JN-RM-0419-21, Jul. 2021.

[11] E. Henneman, “Relation between Size of Neurons and Their Susceptibility to Discharge,”Science (80-.)., vol. 126, p. 1345, 1957.

[12] E. Henneman, H. P. Clamann, J. D. Gillies, and R. D. Skinner, “Rank order of motoneurons within a pool: law of combination,”J. Neurophysiol., vol. 37, pp. 1338–1349, Jul. 1974.

[13] R. B. Stein, E. R. Gossen, and K. E. Jones, “Neuronal variability: noise or part of the signal?,”Nat. Rev. Neurosci., vol. 6, p. 389, 2005.

[14] B. Gustafsson and M. J. Pinter, “On factors determining orderly recruitment of motor units: a role for intrinsic membrane properties,”Trends Neurosci., vol. 8, pp. 431–433, Jul. 1985.

[15] J. E. Desmedt and E. Godaux, “Fast motor units are not preferentially activated in rapid voluntary contractions in man,”Nature, vol. 267, p. 717, 1977.

[16] T. Oya, S. Riek, and A. G. Cresswell, “Recruitment and rate coding organisation for soleus motor units across entire range of voluntary isometric plantar flexions,”J Physiol, vol. 587, pp. 4737–4748, 2009.

[17] E. J. van Zuylen, C. C. Gielen, and J. J. Denier van der Gon, “Coordination and inhomogeneous activation of human arm muscles during isometric torques,”J Neurophysiol, vol. 60, pp. 1523–1548, 1988.

[18] C. K. Thomas, B. H. Ross, and B. Calancie, “Human motor-unit recruitment during isometric contractions and repeated dynamic movements,”J Neurophysiol, vol. 57, pp. 311–324, 1987.

[19] C. K. Thomas, B. H. Ross, and R. B. Stein, “Motor-unit recruitment in human first dorsal interosseous muscle for static contractions in three different directions,”J Neurophysiol, vol. 55, pp. 1017–1029, 1986.

[20] A. W. Monster and H. Chan, “Isometric force production by motor units of extensor digitorum communis muscle in man,”J Neurophysiol, vol. 40, pp. 1432–1443, 1977.

[21] A. Adam and C. J. De Luca, “Recruitment order of motor units in human vastus lateralis muscle is maintained during fatiguing contractions,”J Neurophysiol, vol. 90, pp. 2919–2927, 2003.

[22] B. W. Fling, C. A. Knight, and G. Kamen, “Relationships between motor unit size and recruitment threshold in older adults: implications for size principle,”Exp Brain Res, vol. 197, pp. 125–133, 2009.

[23] J. V Basmajian, “Control and Training of Individual Motor Units,”Science (80-.)., vol. 141, no. 1962, p. 440, 1963.

[24] N. J. Marshall et al., “Flexible neural control of motor units,”bioRxiv, p. 2021.05.05.442653, May 2021.

[25] D. Y. Barsakcioglu, M. Bräcklein, A. Holobar, and D. Farina, “Control of Spinal Motoneurons by Feedback From a Non-Invasive Real-Time Interface,”IEEE Trans. Biomed. Eng., vol. 68, no. 3, pp. 926–935, Mar. 2021.

[26] A. Del Vecchio, A. Holobar, D. Falla, F. Felici, R. M. Enoka, and D. Farina, “Tutorial: Analysis of motor unit discharge characteristics from high-density surface EMG signals,”J. Electromyogr. Kinesiol., vol. 53, p. 102426, Aug. 2020.

[27] M. Bräcklein, J. Ibanez, D. Y. Barsakcioglu, and D. Farina, “Towards human motor augmentation by voluntary decoupling beta activity in the neural drive to muscle and force production,”J. Neural Eng., vol. 18, no. 1, p. 16001, Nov. 2020.

[28] J. Rochon, M. Gondan, and M. Kieser, “To test or not to test: Preliminary assessment of normality when comparing two independent samples,”BMC Med. Res. Methodol., vol. 12, no. 1, pp. 1–11, Jun. 2012.

[29] D. Patikas et al., “Electromyographic changes of agonist and antagonist calf muscles during maximum isometric induced fatigue,”Int. J. Sports Med., vol. 23, no. 4, pp. 285–289, May 2002.

[30] M. D. Binder, R. K. Powers, and C. J. Heckman, “Nonlinear Input-Output Functions of Motoneurons,”Physiology, vol. 35, no. 1, pp. 31–39, Jan. 2020.

[31] C. J. Heckman, M. Johnson, C. Mottram, and J. Schuster, “Persistent inward currents in spinal motoneurons and their influence on human motoneuron firing patterns,”Neuroscientist, vol. 14, no. 3, pp. 264–275, 2008.

[32] M. Gorassini, J. F. Yang, M. Siu, and D. J. Bennett, “Intrinsic Activation of Human Motoneurons: Reduction of Motor Unit Recruitment Thresholds by Repeated Contractions,”J. Neurophysiol., vol. 87, no. 4, pp. 1859–1866, 2002.

[33] J. Ibáñez, A. Del Vecchio, J. C. Rothwell, S. N. Baker, and D. Farina, “Only the Fastest Corticospinal Fibers Contribute to β Corticomuscular Coherence,”J. Neurosci., vol. 41, no. 22, pp. 4867–4879, Jun. 2021.

[34] B. M. ter Haar Romeny, J. J. van der Gon, and C. C. Gielen, “Relation between location of a motor unit in the human biceps brachii and its critical firing levels for different tasks,”Exp Neurol, vol. 85, pp. 631–650, 1984.

[35] H. E. Desmedt and E. Gidaux, “Spinal motoneuron recruitment in man: rank deordering with direction but not with speed of voluntary movement,”Science (80-.)., vol. 214, p. 933, 1981.

[36] A. Nardone, C. Romanò, and M. Schieppati, “Selective recruitment of high-threshold human motor units during voluntary isotonic lengthening of active muscles.,”J. Physiol., vol. 409, no. 1, pp. 451–471, Feb. 1989.

[37] C. M. Laine, E. Martinez-Valdes, D. Falla, F. Mayer, and D. Farina, “Motor Neuron Pools of Synergistic Thigh Muscles Share Most of Their Synaptic Input,”J. Neurosci., vol. 35, no. 35, pp. 12207–12216, 2015.

